# Engineering directional phosphoryl flow enables programmable signaling dynamics in bacteria

**DOI:** 10.1101/2025.11.17.688798

**Authors:** Marcos Nieves, Juan Manuel Valle, Joaquín Dalla Rizza, Federico Carrión, Pablo Naranjo-Meneses, Nicole Larrieux, Ilka B. Bischofs, Alejandro Buschiazzo, Felipe Trajtenberg

## Abstract

Cells must make critical decisions by integrating information from a constantly changing environment to ensure their survival. They rely on intricate signaling networks to detect external and internal cues to trigger specific responses, yet these systems are generally viewed as components wired with simpler linear connections. Bacterial phosphorelay systems offer a versatile framework for studying more complex connections and engineering new biological circuits. Here, using the well-studied *Bacillus subtilis* sporulation phosphorelay, we demonstrate that the directionality of information flow can be reprogrammed to generate different dynamic responses. We show that phosphoryl-transfer reversibility is an evolvable trait encoded in conserved, surface-exposed motifs of two-component system proteins. Unidirectional phosphoryl-transfer generates a short-term information storage mechanism, enabling signal integration over time and allowing phosphatases, acting at different levels, to produce different outcomes. In contrast, a bidirectional system enhanced the action of phosphatase activity early in the pathway. The ability to control phosphoryl-transfer equilibria opens exciting avenues for designing sophisticated synthetic signaling systems with enhanced decision-making capabilities.

## INTRODUCTION

Cells are constantly evaluating their environment and collecting information about their internal status, performing complex biological computations to make decisions. The adaptive response triggered by sensory systems allows for survival under highly fluctuating environments, including cell communication, nutrient limitation, temperature shifts or desiccation. TCSs are widely spread signaling pathways in bacteria and archaea, but also present in fungi and plants [1]. Information travels from a sensory histidine kinase (HK) to an effector response regulator (RR) by means of phosphoryl-transfer reactions between specific histidine and aspartate residues [2].

An interesting feature of TCS chemistry, is that phosphoryl-transfer equilibria can be displaced towards the histidine or the aspartate residues (S1 Fig). Based purely on energetic grounds, the phosphoryl group *en transfer* tends to go to the histidine’s imidazole, and away from the carboxylate side chain of aspartate [3]. However, different mechanisms can overcome this energetic uphill [4, 5], eventually favoring aspartate phosphorylation. Among such mechanisms, the donor:acceptor interatomic distance within the reaction center [4, 5], HK allosteric conformational rearrangements, as well as RR dimerization [4], have all been shown to contribute to determining the overall directionality of the reaction. In the simplest TCS configuration, with a HK and a RR, a strongly unidirectional system is needed to promote turnover and efficient signaling [4].

Phosphorelays are a complex type of TCS, comprising several intermediate proteins between the sensory HK and the effector RR. Longer phosphoryl-transfer cascades as this enables a rich integrative signaling processes that regulate important biological phenomena [6, 7]. The phosphoryl group must travel through multiple reaction centers with disparate directions: His to Asp, as well as Asp to His. This raises several fundamental questions: Are phosphoryl-transfer reactions fully reversible in these pathways? Can the equilibrium of these reactions be engineered to tune their directionality? What properties can emerge in these networks?

To answer these questions, we turned our attention to the sporulation regulatory pathway from *B. subtilis* [6]. Sporulation is an important developmental process that generates a highly specialized cell, the dormant endospore, which sustains a stagnated metabolism that renders them extremely resistant to environmental stress [8]. Several HKs (KinA, KinB, KinC, KinD and KinE) [9] monitor internal and external signals in the *B. subtilis* cells (S2 Fig). This sensory information is channeled through intermediate components, Spo0F and Spo0B, that transfer the phosphoryl moiety to the downstream master regulator Spo0A [6]. Spo0F and Spo0A are RRs, while Spo0B is a phosphotransferase. The phosphorelay is also affected by dedicated phosphatases, such as receptor aspartyl phosphatases (Raps) and Spo0E-like proteins, that drain the phosphoryl group from Spo0F and Spo0A, respectively [10, 11]. Spo0A controls different outcomes, including sporulation [6], biofilm formation [12], cannibalism [13] and cellular competence [14]. Once a certain level of phosphorylated Spo0A is reached [15], a complex phenotypic reprogramming is triggered, ultimately ending with the formation of the endospore [15]. In addition to other downstream target genes, Spo0A is also involved in regulating the expression of components of the phosphorelay pathway itself, thus shaping complex behaviours under different conditions [16]. The *B. subtilis* sporulation phosphorelay is very well studied both experimentally and computationally [14, 17, 18], an ideal model to address our questions.

Here we show that the sporulation phosphorelay from *B. subtilis* can be reprogrammed by specific mutations at key protein:protein interfaces, reshaping the output dynamics of the signaling system. We demonstrate that natural phosphoryl-transfer equilibria are displaced towards downstream components in each reaction pair along the sporulation path. A defined His-Asp distance is determined by conserved motifs on the two RRs that interact with Spo0B. These interface contacts position each RR in a different pose, suggesting that directionality determinants are encoded within the RR structure and they are evolutionarily selected. Through experimental approaches and mathematical modelling, we show that the *B. subtilis* sporulation phosphorelay can transiently store information in the form of phosphorylated Spo0A, enabling a rheostat-like behavior of the system depending on the kinase/phosphatase activity ratio. In contrast, imposing a reversible phosphoryl-transfer equilibrium enhances the action of Rap phosphatases. These findings provide a new understanding of how protein-level mechanisms can shape network-wide signaling dynamics and emergent properties.

## RESULTS

### Unidirectional phosphoryl-transfer towards downstream phosphorelay components

Considering that 3D structures of the Spo0F:Spo0B complex from *B. subtilis* have been known for decades, to better understand the phosphorelay’s signaling mechanism we determined the crystal structure of the Spo0B:Spo0A binary complex. Its structure was solved in two different crystal forms (S1 Table) at 2.1 and 3.5 Å resolution. The refined atomic model shows the expected 2:2 phosphotransferase:RR stoichiometry (Fig 1A), similarly to Spo0F:Spo0B. The transferase interacts with Spo0A mainly through helices α1 and α2 on the DHp (***D***imerization and ***H***istidine ***p***hosphotransfer) domain with the α1α5 face of Spo0A, analogous to many other HPt:RR (HPt stands for ***H***istidine ***P***hospho***t***ransferase) and HK:RR complexes [5, 19–23]. In each of the independently refined Spo0B:Spo0A complexes (7 in total), one of the two Spo0A molecules displays better defined electron density, resulting in lower refined B factors. Interestingly, these can be explained by additional contacts established between Spo0A and the pseudo-ATP-binding domain (pseudoABD) of Spo0B, breaking the symmetric structure of Spo0B.

**Fig. 1.**
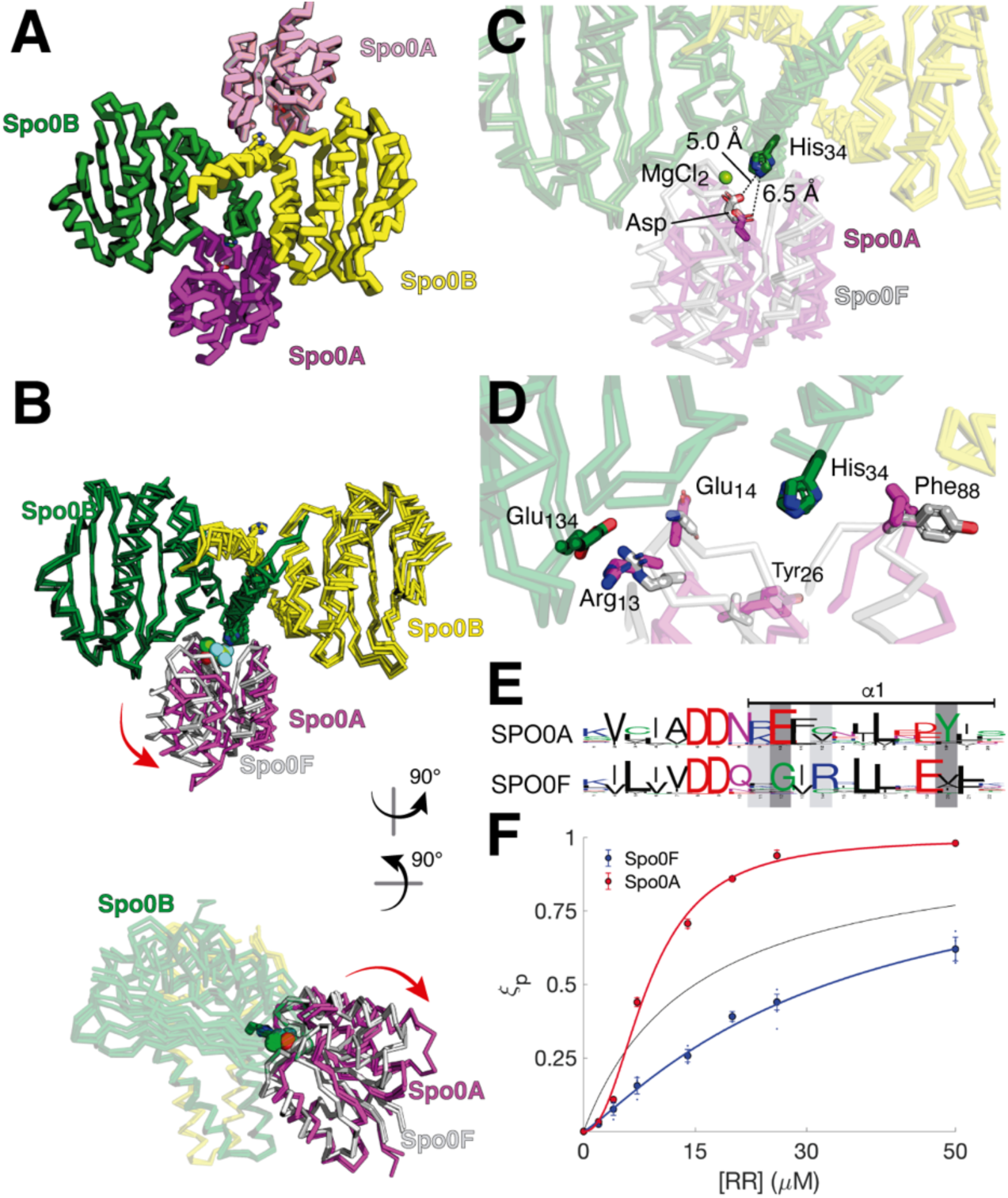
Crystal structure of the Spo0B:Spo0A complex. A) Ribbon representation of the complex. Two protomers of Spo0B (depicted in green and yellow) interact with two receiver domains of Spo0A (pink and magenta). B) Structural alignment of Spo0B:Spo0A and Spo0F:Spo0B complexes, using the RR-binding site of Spo0B as reference (Spo0B residues 30 to 67). Spo0F is shown as white ribbon. Red arrows indicate the composite movement that relates the position of Spo0F and Spo0A in the complexes. C) Close-up of the active sites showing the distance between His and Asp within the complexes. The distance between the reactive His and Asp in the Spo0B:Spo0A complex is 6.6 Å ± 0.2 (standard deviation estimated using per atom-corrected DPI [51]), whereas in the Spo0F complex the distance is 5.0 ± 0.5 Å. D) Protein:protein interacting surfaces between Spo0B and Spo0F (white) or Spo0A (magenta) is shown. Selected set of residue pair interactions are depicted as sticks. E) Sequence conservation logo of the β1 α1 loop and helix α1 region of Spo0F and Spo0A. F) The degree of phosphorylation (ξp) was calculated as the relative amount of RR∼P over all phosphorylated proteins (RR∼p+Spo0B∼P) as obtained by densitometric quantification of phosphotransfer titration assays of Spo0F (blue) and Spo0A (red), respectively. Circles show mean and standard deviation from three independent experiments. Lines represent fits of the data to a Hill function. Green dashed lines indicate the reactions in equimolar conditions. The thin black line represents the expected behavior in a fully reversible equilibrium.

Despite overall similar organizations of Spo0A and Spo0F with respect to Spo0B, the superposition of each complex using Spo0B’s α1/α2 helices as reference, reveals a significant shift in the position of Spo0A, involving translation and rotation relative to Spo0B (Fig 1B). As a result, the phosphorylatable Asp of Spo0A is placed significantly farther away from Spo0B’s His (6.6 Å) compared to the equivalent Asp in Spo0F (5.00 Å), as shown in Fig. 1C. Interestingly, the Spo0F:Spo0B complex exhibits a larger REC:pseudoABD interface, which buries a surface of ∼300 Å^2^ and includes hydrogen-bonds and van der Waals contacts. In contrast, the equivalent Spo0B:Spo0A interface totals ∼90 Å^2^ with only a few van der Waals interactions (Fig 1D). From a structural point of view, the receiver domain of Spo0A and Spo0F are very similar (Fig 1E). However, both complexes show specific sets of contacts with Spo0B that are differential and engage evolutionarily conserved residues [24]. Notably, key differences lie on the DHp:REC interface. The highly conserved motif REXX (including Arg18 and Glu19 at the beginning of helix α1) in Spo0A, is substituted to XGXR in Spo0F (Fig 1E and S3). This effectively moves the position of the Arg by one helix turn (Arg16_Spo0F_ → Arg19_Spo0A_), while introducing a bulkier adjacent residue (Gly14_Spo0F_ → Glu18_Spo0A_). In addition, the conserved Phe92 from Spo0A is packed against Spo0B residues Met37 from α1, and Val63, Ala65 and Lys67 from α2. On the other hand, Spo0F includes a Tyr in the same position, which is solvent exposed. Taking all together, the data imply an evolutionary pressure to correctly place each RR protein with respect to the Spo0B partner, placing the phosphorylatable His and Asp residues at specific and distinct distances. A longer interatomic distance between phosphoryl-donor and -acceptor are known to correlate with a stronger directionality from P∼His to Asp [4]. This suggests that the Spo0F:Spo0B and Spo0B:Spo0A phosphotransfer equilibriums could be differentially displaced towards either the His or the Asp, respectively.

To confirm our prediction, we studied the reaction of the phosphorylated phosphotransferase Spo0B (P∼Spo0B) with the two RR *in vitro* using radiolabeled ATP[γ-^33^P]. A fixed concentration of P∼Spo0B [15 µM] was exposed to varying concentrations of Spo0A (or Spo0F) from 0 to 50 µM. Equilibrium was reached after 1 min. The distribution of phosphate among the two proteins was analyzed by autoradiography and showing an accumulation of phosphorylated RR [RR∼P] with a concomitant decrease of Spo0B∼P as the concentration of RR increases (S4 Fig). The fraction of phosphate on the RR over all phosphorylated proteins (ξ_p_, defined as [RR∼P]/([RR∼P]+[Spo0B∼P])) for both Spo0A (red symbols) and Spo0F (blue symbols) was well described by a Hill function (Fig 1F). The Spo0B-Spo0F phosphoryl-transfer showed low levels of cooperativity (Hill coefficient n = 1.2 for Spo0F) while the reaction between Spo0B and SpoA showed strong cooperativity (n = 2.2). Moreover, when compared to the equilibrium distribution of an ideal fully reversible phosphoryl-transfer reaction (black line), it is clear that phosphoryl-transfer between Spo0B and Spo0F was shifted towards the phosphotransferase (Asp to His direction), while the phosphoryl-transfer between Spo0B and Spo0A is shifted towards the RR (His to Asp direction). Together these findings indicate that the *B. subtilis* sporulation phosphorelay is configured to facilitate a directional phosphoryl-flux from upstream to downstream signaling components.

### Reconfiguring reversibility in TCS phosphorelays

Next, we tested whether we could modulate the phosphoryl-transfer directionality by mutating residues in the binding interface of Spo0B with Spo0F and Spo0A. Informed by the structure of the complexes, we chose eleven Spo0F (F1-11) and three Spo0A (A1-3) variants by grafting five different interface motifs (S5 Fig). For example, Spo0F variants F1 and F2 involve grafting of one or more residues from motif 1 of Spo0A to Spo0F, F3 and F4 are modifications at motif 3, and F5 to F7 are combinations of modifications in motifs 1 and 3 (see full details in S2 Table and S5 Fig). For Spo0A, we included the double mutant Y108A and H122A that interferes with dimerization (A1), a point mutant E14G (A2), which is part of the conserved motif interacting with Spo0B and a truncated variant on Spo0A’s, including only the receiver domain (A3).

For all proteins variants we measured the distribution of phosphoryl-groups after incubation with P∼Spo0B as before (Fig. 2A). For Spo0F (blue bars), substitutions at motif 3 (present in F3, F4, F6, F7 and F8) resulted in a significant shift of the phosphoryl-equilibria towards the RR compared to wild-type Spo0F, losing directionality. The strongest effects were seen for F3 and F4, both containing a K56I substitution that is located two residues after the phosphorylatable Asp. In contrast, mutant F9, which includes substitutions at motifs 1, 2 and 4, shifted the reaction towards phosphorylation of the phosphotransferase Spo0B. This effect was lost by including K56I and E86Q substitutions on motif 3 (F10 and F11), suggesting a compensatory effect between motifs. Notably, all three Spo0A mutants shifted the phosphoryl-transfer equilibrium towards Spo0B (red bars).

**Fig. 2.**
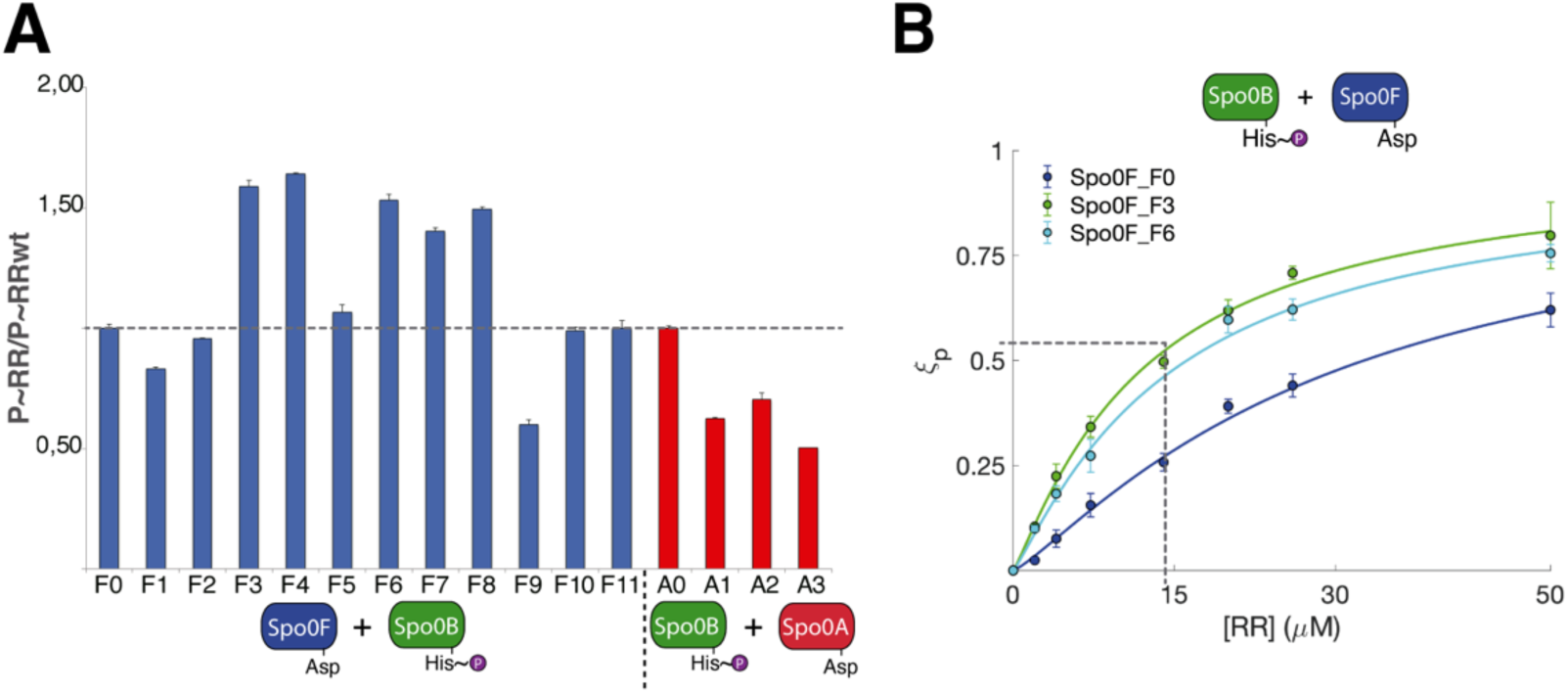
Mutant grafting and phosphotransfer directionality analyses. A) Level of phosphorylation of RR variants (20 µM) after reaching equilibrium with P∼Spo0B (15 µM), normalized against wild type Spo0F (F0) or Spo0A (A0). Reactions were analyzed by autoradiography. Spo0F wild type (F0) and its variants are shown in blue, whereas Spo0A wild type (A0) and its variants are depicted in red. B) Phosphotransfer equilibrium titration experiments for wild type Spo0F (F0) and its variants F3 and F6. P∼Spo0B (15µM) was incubated with varying concentrations of the RR (from 0 to 50 µM) and analyzed after reaching equilibrium. Phosphorylation levels were determined (circles) and Hill functions were fitted to the data obtained (solid lines).

We next performed titration assays with two Spo0F variants F3 (green line) and F6 (cyan line) and compared it to wild type Spo0F (F0, blue line) as shown in Fig 2B. At equimolar concentrations of Spo0B and Spo0F, the phosphoryl-group were almost evenly split between both proteins (dashed lines), indicative of a fully reversible phosphoryl-transfer equilibrium. Interestingly, a single substitution K56I on Spo0F (F3) was enough to make the phosphoryl-transfer fully reversible

Taken together these data show that directionality of both phosphotransferase reactions can be shifted from the directional phosphoryl-transfer, as observed in wild type, to a fully reversible equilibrium.

### Phosphoryl flow directionality is essential for phosphorylation-dependent short-term memory

Collectively, our results suggest that phosphoryl-transfer directionality in the sporulation phosphorelay is a trait that can be accessed by evolution. We thus wondered whether the directionality of phosphoryl-transfer reactions could confer specific properties to a signaling system. The overall organization of the sporulation regulation pathway (S2 Fig) led us to speculate that phosphoryl-transfer reversibility could affect the action of phosphatases at the level of Spo0F. To investigate this possibility, we used a deterministic mass-action kinetic model of the signaling system (Fig 3A) based on reported data [18, 25] . The kinetic model included KinA, Spo0F, Spo0B, Spo0A and RapA. Feed-forward loops (in KinA, Spo0F, Spo0A and RapA) were modelled as Hill equations, dependent on the levels of P∼Spo0A (full details in Material and Methods). Under a regime of unidirectional phosphoryl-transfer reactions, as observed in our biochemical characterizations (Fig 1F), kinetic modelling indicated that even at very high concentration of phosphatase activity, information accumulates over time (Fig 3B, left panel), in the form of total amount of P∼Spo0A.

**Fig. 3:**
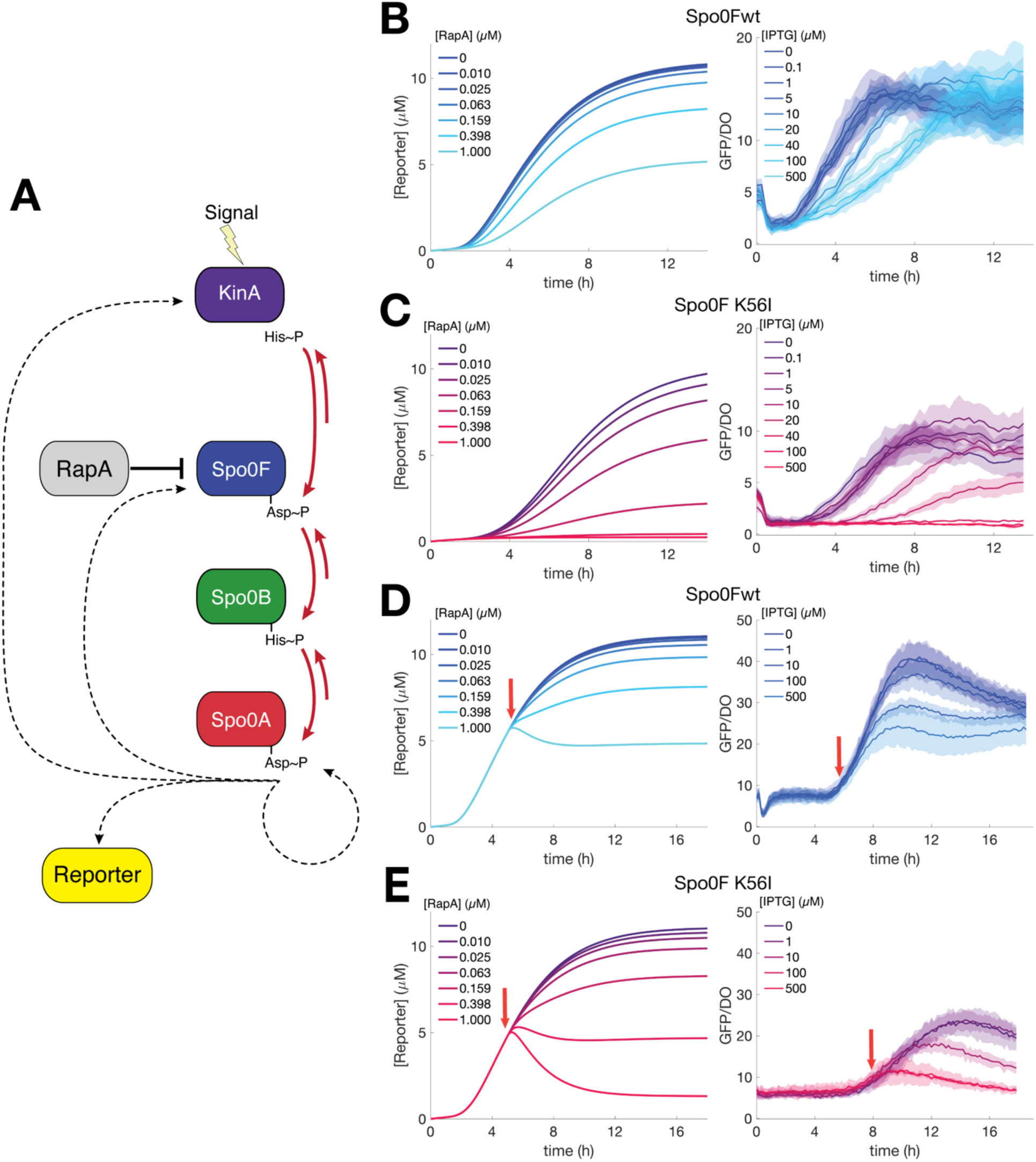
Directionality in phosphoryl-transfer affects signal integration and information storage. A) Schematic representation of the ordinary differential equations included in the kinetic model. Red arrows indicate phosphoryl-transfer reactions. Dashed arrows depict transcriptional activation and are modelled as an activating Hill function dependent on P∼Spo0A concentration. B) Left panel shows the simulations of the sporulation system under activating conditions with wild type spo0F and under different steady state RapA concentrations. The reporter concentration is plotted as a function of time. The parameters that describe the phosphotransfer reactions were approximated to describe the observed displacements of the equilibria according to the titration experiments. Right panel are the time courses of sporulation assays in engineered B. subtilis strains expressing wild type Spo0F (strain BSS117). Sporulation was induced by incubating cells in poor media under different IPTG concentrations (setting distinct RapA intracellular levels). Thick lines represent the mean GFP/DO ratio, and the shaded region indicates the one standard deviation. C) Similar to B but using Spo0F_K56I variant (left simulations and right the sporulation assay with strain BSS118). Note that reporter levels in strain carrying wild type Spo0F increase even at the highest IPTG concentrations, while in these conditions, the pathway remains turned off in the strain with spo0F_K56I mutant. D) Similar to B, but RapA was induced (indicated with red arrows) after the system showed signs of activation. In the right panel IPTG was added at different concentrations after approximately 5 hours of incubation. E) Similar to D but using the BSS118 strain with the spo0F_K56I mutant.

To test the predictions derived from the kinetic model, we constructed *B. subtilis* strains that included a super folder green fluorescent protein (GFP) reporter gene under the *spoIIE* high-threshold promoter. This was generated with a *spo0F*, *spo0E* and *rapA* triple knockout strain. RapA was inserted ectopically under an IPTG-inducible *Phyperspank* promoter, to enable the control of phosphatase expression in the absence of the PhrA inhibitor. On this GFP reporter strain, we then reverted the *spo0F* knockout by homologous recombination at the endogenous locus with wild type *spo0F* (strain BSS117). Exponentially growing *B. subtilis* cells were switched from a rich to a poor medium to induce sporulation at 37 °C, and fluorescence was measured over time. Different IPTG concentrations were used to control RapA expression at different steady state levels. The back-complemented strain showed an increase in fluorescence over time, even at the highest IPTG concentrations (Fig 3B, right panel). Although higher RapA levels slowed down the activation process, eventually cells reached the maximum signal (Fig 3B, right panel).

On the other hand, simulations of a phosphorelay with a fully reversible Spo0F:Spo0B phosphoryl-transfer step, as measured for the Spo0F-K56I mutant (F3, Fig 2B), showed that a high enough RapA concentration prevents accumulation of P∼Spo0A and, therefore, precludes the activation of high-threshold promoters (Fig 3C, left panel). Note that, by including the Spo0F_K56I variant, only the Spo0F:Spo0B step is being modified, such that the pathway retains some directionality towards Spo0A phosphorylation, albeit to a lesser extent. As predicted by the model, the back-complemented strain with Spo0F-K56I variant was unable to activate the Spo0A-controlled pathway at the highest RapA concentrations (Fig 3C, left panel). These results indicate that phosphoryl-transfer reversibility is relevant, and it is configured in such a way that the sporulation regulation system can slowly accumulate information over time, in the form of P∼Spo0A, even under high RapA phosphatase activity.

In a different experimental setup, where RapA activity is induced after the system was activated, the signaling pathway with wild type Spo0F stored information in the form of P∼Spo0A (Fig 3D). In contrast, the *spo0F-K56I* strain showed a complete reset of the system at high RapA concentrations (Fig 3E), indicating that pathway activation can be reversed, consistent with our simulations. Our kinetic model further predicts that, unlike the observed effect of RapA phosphatase in the *spo0F* wild-type background (Fig. 3B), phosphatase activity acting at the Spo0A level should reset the system (Fig. S6A). This prediction was confirmed experimentally by monitoring *spoIIE* promoter activity in a *spo0E rapA* double knockout strain, in which Spo0E was expressed under IPTG control (S6B Fig). Altogether, these results indicate that the directionality of phosphoryl flow in the sporulation phosphorelay is organized to transiently record and store information, effectively functioning as a short-term memory system that can be reset by downstream phosphatase activity.

## DISCUSSION

This study reveals that the directionality of phosphoryl-transfer reactions plays a pivotal role in information storage, signal integration and shaping of output responses. Challenging a common view of signaling networks as simple and linear pathways, phosphorelays offers the opportunity to create more complex network connections. We now show that by modifying conserved, surface-exposed motifs, we can engineer networks with distinct phosphoryl-transfer reversibility, highlighting how the inherent biochemistry of protein-protein interactions can be manipulated to generate sophisticated, non-intuitive network behaviors.

The phosphotransfer reaction between Spo0F and Spo0B is shifted towards histidine phosphorylation, as expected on energetic grounds. However, the opposite is observed in the Spo0B:Spo0A phosphotransfer equilibrium, indicating that each RR protein encodes the directionality determinants in its 3D structure. The crystal structure of the Spo0B:Spo0A complex revealed a longer His-Asp distance compared to that of Spo0F:Spo0B. The distance between the phosphoryl donor and acceptor residues has been proposed to be a key directionality determinant [4, 5]. Spo0A dimerization is also a directionality determinant, a feature shared with other RRs [4]. Thus, RR dimerization not only triggers an effector response, but in addition plays a role in minimizing back-transfer of the phosphoryl group to the HK, and thereby, maximizes a unidirectional, downstream phosphoryl flux.

The structural comparison between Spo0B:Spo0A and Spo0F:Spo0B uncovered specific interfaces that seem key for the proper positioning of the RR within the complex. We had previously shown that HK:RR interactions are “slippery” [5] and that such plasticity allows for the relocation of the RR within the complex guided by interactions with additional interfaces, such as those between the ABD and REC domains [4]. In the case of the TCS DesK:DesR from *B. subtilis*, the ABD:REC contacts shift the position of the RR, bringing the phosphorylated Asp residue closer to the conserved Gln [4]. Analogously, we now reveal that the interactions between the pseudoABD domain of Spo0B and Spo0F are tighter than with Spo0A. In addition, Spo0F and Spo0A show conserved differences in positions that are directly involved in the DHp:REC interaction. These two traits, end up placing Spo0F in a shifted position with respect to Spo0A (Fig 1) in complex with Spo0B, resulting in a shorter His-Asp distance, and suggesting that phosphotransfer directionality is encoded in the protein:protein interaction surfaces. Sequence conservation indicates that directionality might also be an evolvable property, a matter for future studies. The fact that directionality determinants between Spo0F and Spo0A can be grafted and exchanged, indeed suggests that directionality is transferrable, and thus evolvable under selective pressure.

Notably, the identified *spo0F_K56I* mutant, which shifts the phosphoryl-transfer equilibrium, allowed us to assess the relevance of phosphorelays with different directional transmission of information. As we now show, unidirectional phosphoryl-transfer in the sporulation pathway allows a mechanism of short-term memory, whereby P∼Spo0A can be accumulated over time. This stored information is reset by the action of Spo0A-specific phosphatases like Spo0E (S6 Fig). All things considered, a conceptual model implies a system that can switch between recording, storing and resetting phases (Fig 4). Moreover, the system can work as a rheostat, offering a gradual and controlled response to changes in kinase/phosphatase ratios (S7 Fig). This rheostat-like behavior is enabled by the phosphoryl-transfer unidirectionality and means that rather than simply turning “on” or “off,” the pathway can fine-tune its activation dynamics across a continuous spectrum. This allows it to effectively integrate the diverse signals from various sensor HKs and phosphatases.

**Figure 4.**
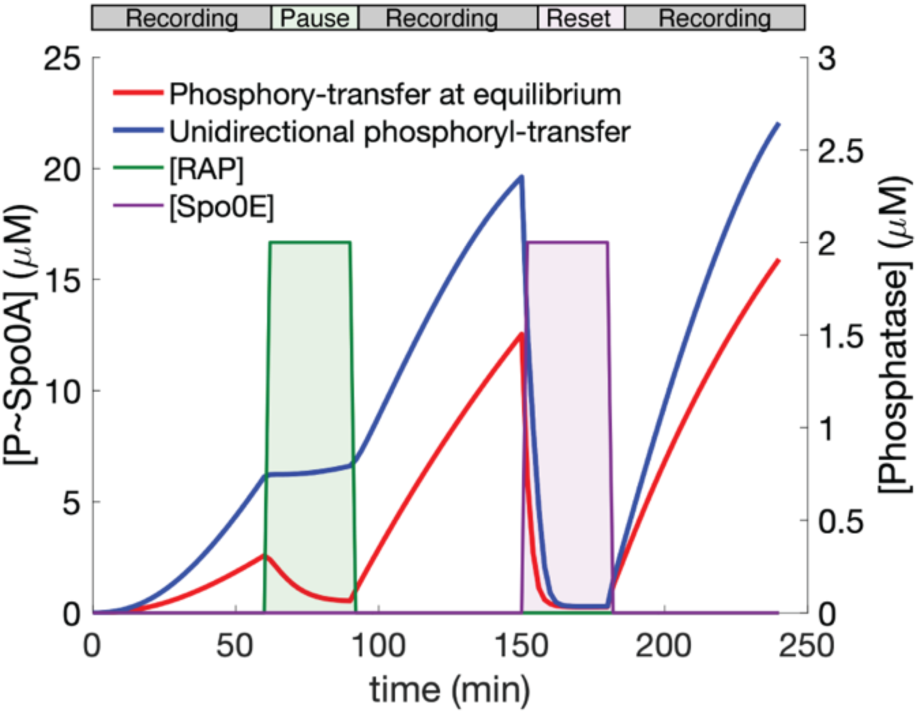
Information recording, storage and reset in the B. subtilis sporulation regulation pathway. P∼Spo0A accumulation time courses obtained by kinetic simulations in systems with fully reversible equilibria (red line) or with unidirectional phosphoryl-transfer transmission (blue line). During the first 60 min, the reactions proceed in the absence of phosphatases, with KinA activity recording information as P∼Spo0A. When a pulse of RapA phosphatase (green bar) was included, the information was stored only in the unidirectional system, establishing a short-term memory. A pulse of Spo0E phosphatase (magenta bar), after a second recording phase, resulted in a reset of the information.

The sporulation pathway exhibits distinct behaviors under varying growth conditions. In some cases, it follows a pulsatile pattern [18, 26], while in others, cells gradually accumulate P∼Spo0A until reaching a specific threshold that triggers pathway activation [15, 27]. The promoter controlling *rapA* can also show pulsatile dynamics [28]. Such different dynamics could be generated due to different kinase/phosphatase ratios, as well as by triggering alternative dedicated phosphatase activities, like RapA or Spo0E. However, further studies are needed in less-engineered strains and in different culture scenarios to fully understand how *B. subtilis* makes the decision to sporulate.

A striking aspect of the sporulation pathway is the ability to generate different cellular outcomes while integrating a myriad of signals [29]. This has been further supported by recent observations showing that different HKs can trigger alternate cell fates [30]. The presence of different Spo0A promoters with distinct thresholds [16] could imply, not only the ability to control sequential steps along the sporulation program, but also the capacity to trigger alternate genetic programs due to different activation kinetics. The latter, as we see in our study, are coupled to time-sensitive activation or repression of phosphatases in the sporulation phosphorelay. The modularity and signal-integrating features of this pathway represent a blueprint for building complex, multi-input genetic circuits with tailored behaviors.

The significance of phosphoryl-transfer reversibility extends beyond the *B. subtilis* sporulation pathway. The chemical versatility of His to Asp phosphotransfer allows generating reactions with varying degree of reversibility (S1 Fig). Phosphotransfer directionality has been proposed to warrant efficient signal transmission [4], and also to allow for “phosphate sinks” in several phosphorelay systems [31–33]. In summary, we have shown that phosphotransfer directionality is a key determinant to shape how information is integrated and processed. We also demonstrated that directionality can be reprogrammed and can thus be subject to evolutionary pressure. This adds a completely new layer of complexity in rewiring signaling pathways, involving not only the inter-protein specificity determinants [34], but also the directionality of the phosphoryl-transfer reactions. Engineering signaling pathways could benefit from the ability to modulate the reversibility of such equilibria in order to generate more efficient and tunable information-processing systems for Synthetic Biology.

## MATERIALS AND METHODS

### Cloning and protein purification

Cloning and protein purification were performed as described previously [35]. The plasmid for recombinant production of KinA (pETM_KinA) was constructed by cloning into a pETM-14 plasmid (kindly provided by Frank Lehman) using primers KinA_F and KinA_R by restriction-free cloning [36]. Similar procedure was followed to generate plasmids pQE80_Spo0A and pQE80_Spo0A_REC, using primers Spo0A_F, Spo0A_R and Spo0A_REC_R. The primers spo0E_up_F, spo0E_up_R and Spo0E_down_F, spo0E_down_R were used to amplify 500-bp fragments upstream and downstream of spo0E, respectively. The DNA fragments were fused by PCR and cloned into the pMAD vector by RELC using the enzymes SalI and BglII. Plasmids pQE80_Spo0F, pQE80_Spo0B, pET_Spo0B_strep, and all the Spo0F and Spo0A mutants tested, were obtained by synthesis and subcloning services (Genscript). Tables of strains, plasmids and oligonucleotide primers are included as supplementary data (S3-S5 Tables).

Recombinant proteins were expressed as N-terminally His6-tagged fusions in *E. coli* strain BL21 (DE3) pLysS as previously performed for other TCS proteins [37]. Transformed bacteria were grown in TB medium, and recombinant protein expression was induced during exponential growth by adding 1 mM IPTG for 18 hours at 20 °C. Cells were harvested by centrifugation and resuspended in lysis buffer (50 mM Tris-HCl pH 8.0, 0.5 M NaCl, 1 mg/mL lysozyme, 0.01% Triton X-100, 20 mM imidazole). Cell lysis was performed using either sonication or an Emulsiflex homogenizer. The lysates were centrifuged, and the soluble fraction was loaded onto a 5 mL HisTrap HP column (Cytiva) equilibrated with buffer A (50 mM Tris-HCl, pH 8.0, 500 mM NaCl, 20 mM imidazole) for IMAC purification. After this initial step, the His6-tag was removed by proteolysis with TEVdt protease [38]. Imidazole from the IMAC step was removed by dialysis against 50 mM Tris-HCl pH 8.0, 0.5 M NaCl. Undigested protein and protease were removed by reverse IMAC. The final purification step was size exclusion chromatography using a HiLoad 16/60 Superdex 75 prep grade column (Cytiva), equilibrated with 20 mM Tris-HCl pH 8.0, 0.1 M NaCl (SEC buffer). All proteins were concentrated to ∼10 mg/mL and stored at -80°C.

### Protein crystallization, data collection and model building/refinement

The crystals of Spo0B:Spo0A complex were obtained by sitting-drop vapor-diffusion at 293 K in 96 well plates (Greiner) assisted by a Honeybee 963 robotic station (Digilab). The protein complex was prepared by mixing purified and concentrated Spo0B (57 mg/mL) and Spo0A-REC (50 mg/mL) at 1:1.1 equivalents. The complex was prepared in the presence of 10 mM MgCl_2_ and diluted in SEC buffer to a final concentration of 25 mg/mL. Drops were set up with 300 nL of protein solution and 300 nL of reservoir, equilibrated against 100 µL of reservoir solution (0.2 M NaF, 20% (w/v) polyethylene glycol 3350). Crystals were cryoprotected by quick-soaking in a solution containing 0.155 M NaF, 22% (w/v) PEG 3350, 25% (v/v) glycerol, and flash frozen in liquid nitrogen.

Single crystal X-ray diffraction data collection was performed at 100 K using either a copper rotating anode home source (Protein Crystallography Facility, Institut Pasteur de Montevideo), or synchrotron radiation at Beamline I-03 (Diamond Light Source, UK). Data processing was carried out with Xia2 [39] or autoPROC [40]. The Spo0B:Spo0A complex was solved by molecular replacement [41] using the Spo0B dimer (PDB ID 1IXM) and the Spo0A receiver domain (PDB ID 1QMP) as search probes. Model rebuilding, refinement and validation were carried out using Coot [42], Buster [43], Phenix [44] and MolProbity tools [45]. Protein visualization, structural analyses, and figure rendering were performed with Pymol (Schrodinger, LLC, 2015). The software used for data processing, structure determination, and analysis was provided by the SBGrid Consortium [46].

### Phosphotransferase assays

Phosphotransfer reactions were performed with P∼Spo0B radiolabeled with ^33^P and different Spo0F and Spo0A variants. P∼Spo0B was prepared by incubating a His-tagged Spo0B with KinA and Spo0F in the presence of 150 μCi [γ-^33^P]-ATP (10 mCi/mL, American Radiolabelled Chemicals, Inc.), unlabeled ATP (400 µM final concentration) and MgCl_2_ for 30 min at room temperature in reaction buffer (100 mM NaCl, 20 mM Tris-HCl pH 8 and 5 mM MgCl_2_). P∼Spo0B was then purified using a Histrap column (Cytiva), concentrated and buffer-exchanged to reaction buffer using ultrafiltration. Proteins were analyzed by SDS-PAGE, quantified, aliquoted and stored at -20°C. Evaluation of the phosphoryl-transfer equilibria was done by incubating 15 µM P∼Spo0B with 20 µM of each of the RRs (Spo0F and Spo0A variants), for 10 min at 25 °C. Reactions were quenched with SDS-PAGE sample buffer and subsequently ran in polyacrylamide gels (Cytiva). Gels were exposed to phosphor screen (for 1 h) and imaged in a Typhoon FLA 7000 imaging system (GE Healthcare). Band intensities were analyzed by densitometry with ImageJ [47] and plotted in Matlab R2020a (MathWorks Inc.). Titration assays were performed as above but with varying concentrations of RRs (from 0 to 50 µM). Each reaction was performed in triplicate.

Phosphotransfer assays between P∼Spo0F and Spo0A, with limiting amounts of Spo0B, were performed following similar procedures as previously [4]. Briefly, Spo0F was phosphorylated with KinA in the presence of ATP and MgCl_2_, and then purified and buffer-exchanged in a Superdex75 10/300 column (Cytiva) equilibrated with reaction buffer. Equimolar amounts of P∼Spo0F and Spo0A (15 µM) were incubated in the presence of 0.3 µM Spo0B. At different time points, the reactions were stopped by adding SDS-PAGE sample buffer and loaded in a Phostag SDS-PAGE. Coomassie blue-stained gels were scanned, and quantification was done by densitometry using ImageJ.

### B. subtilis strain construction

All strains are derivatives of *B. subtilis* 168 1A700 and derived from strain BIB415 (*ΔrapA ΔphrA Δspo0F* [48] by a series of genetic modifications. First, *spo0E* was deleted with the help of a pMAD_spo0E plasmid constructed with primers ST94-99 following established protocols [48]. Next, *rapA* was placed under the control of the P*hyperspank* promoter by transformation with pDR111_rapA [49]. Subsequently, the P*spoIIE*_GFP and Pspo0F_mCherry constructs were inserted into the *yhdG_yhdH* and *bglS* loci, respectively. Finally, the *spo0F* knockout was reverted to either the wild-type allele or the *spo0F-K56I* mutant using plasmids pMAD_Spo0F [48] and pMAD_Spo0F-K56I, obtaining strains BSS117 *ΔrapAΔphrA Δspo0E amyE::[specR Phyperspank_rapA] yhdG_yhdH::[phleoR Psdp_mCherry] bglS::[kanR PspoIIE_gfp]* and BSS118 *ΔrapAΔphrA Δspo0E amyE::[specR Phyperspank_rapA] yhdG_yhdH::[phleoR Psdp_mCherry] bglS::[kanR PspoIIE_gfp] spo0F-K56I*). Similarly, starting with the *spo0E* knockout of strain BIB415, *spo0E* was reinserted in the genome under the control of the P*hyperspank* promoter by transformation with pDR111_spo0E. Subsequently, reporter constructs insertion and *spo0F* knockout restoration were performed identically as for the previous two strains. Thus, obtaining strain BSS119 *ΔrapAΔphrA Δspo0E amyE::[specR Phyperspank_spo0E] yhdG_yhdH::[phleoR Psdp_mCherry] bglS::[kanR PspoIIE_gfp]*. All strains were confirmed by PCR analysis using primers flanking the integration sites, followed by sequencing to verify accurate integration. The resulting strains were stored at -80°C in 15% glycerol until further use.

### Sporulation assay

For sporulation assays under constant RapA and Spo0E levels, strains were grown in LB medium with kanamycin (10 µg/mL) and IPTG (1 mM) overnight at 30°C with shaking at 200 rpm. Fresh cultures were started at an OD_600_ of 0.015 by dilution in LB medium with kanamycin (10 µg/mL) and different concentrations of IPTG (from 0 to 500 µM). Cultures were grown for 3 h at 37°C and 220 rpm, or until they reached an OD_600_ of 0.3. Then, cultures were washed twice in buffer Tris-HCl (10 mM pH 7.4) and resuspended in sporulation medium (SM) with corresponding IPTG concentrations and at a final OD_600_ of 0.1. For each strain and IPTG concentration, 200 µL of each condition was loaded in a 96-well plate with black walls and a clear bottom. Cells were incubated and monitored in a Varioskan (Thermo) with a regime of 30 s of shaking every 4.5 min at 480 rpm, 37°C. Fluorescence (λ_ex_ = 481 nm, λ_em_ = 507 nm) and OD (450 nm) were continuously monitored. Assays were performed in triplicate.

For assays with either RapA or Spo0E induction after sporulation pathway activation, a similar assay was conducted in the absence of IPTG. For this, 180 µL of cultures at an OD of 0.1 in SM medium were inoculated into the 96-well plate, then incubated and monitored using the Varioskan until GFP signal activation was detected and incubated for 30 min. Then, phosphatase expression was induced by adding 20 µL of SM medium with different IPTG concentrations (10-fold relative to the desired final concentration) and cells were monitored under the same conditions.

### Kinetic model and model fitting

Kinetic models were constructed as a set of differential equations based on mass action kinetics using custom scripts in Matlab R2020a (MathWorks Inc.). The model equations were solved numerically using the ode15s solver. Parameters were globally optimized by minimizing the sum of squared residuals (SSRs) of the dataset with the simplex search method as implemented in fminsearch with boundaries [50].

The kinetic model of the sporulation pathway of figure 3 assumes KinA as constitutively active HK. Additional modulators, like Sda, or yet other dedicated phosphatases besides RapA or Spo0E, were not included. We considered cellular quantification and dynamics measured by Eswaramoorthy et al. to set the starting levels and the ratio between synthesis and degradation for each phosphorelay component [25]. Kinetic constants describing phosphoryl-transfer reactions and binding between proteins were taken from Narula et al. [18] and fitting kinetic models to phosphoryl-transfer titration experimental data (S8 Fig). All parameters are publicly available in a Zenodo repository under the following DOI: https://doi.org/10.5281/zenodo.15869743. Reporter expression in Fig. 3 and 4 was modelled using an activation Hill function dependent on P∼Spo0A concentration.

## Acknowledgments

The authors thank Dayana Benchoam and Victor de Lorenzo for helpful discussions and comments. We thank Stephanie Trauth for cloning the pMAD_Spo0E plasmid as part of project ERC-StG 260860. We acknowledge computational and storage services (Maestro cluster) provided by the Institut Pasteur (Paris). AB and FT are associated researchers from PEDECIBA (Programa de Desarrollo de las Ciencias Básicas). IBB is a member of the DFG priority program SPP2389 (BI1213/6-1) and the Marburg cluster of excellence M4C.

## Funding

Agencia Nacional de Investigación e Innovación grants FCE_1_2017_1_13629 and FCE_1_2021_1_166888

DAAD fellowship

Max Planck Society.

## Data and materials availability

Code and models used to generate Figure 3 are publicly available in the Zenodo repository under the following DOI: https://doi.org/10.5281/zenodo.15869743. All data are available in the main text or the supplementary materials.

---

**Fig. S1.**
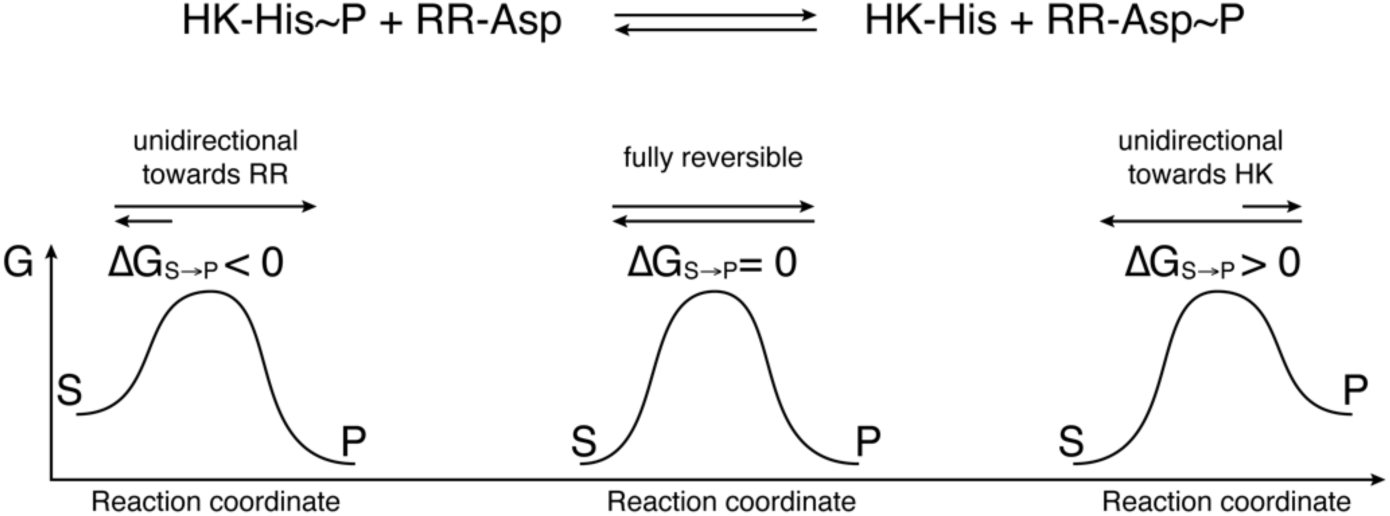
TCS phosphoryl-transfer directionality

**Fig. S2:**
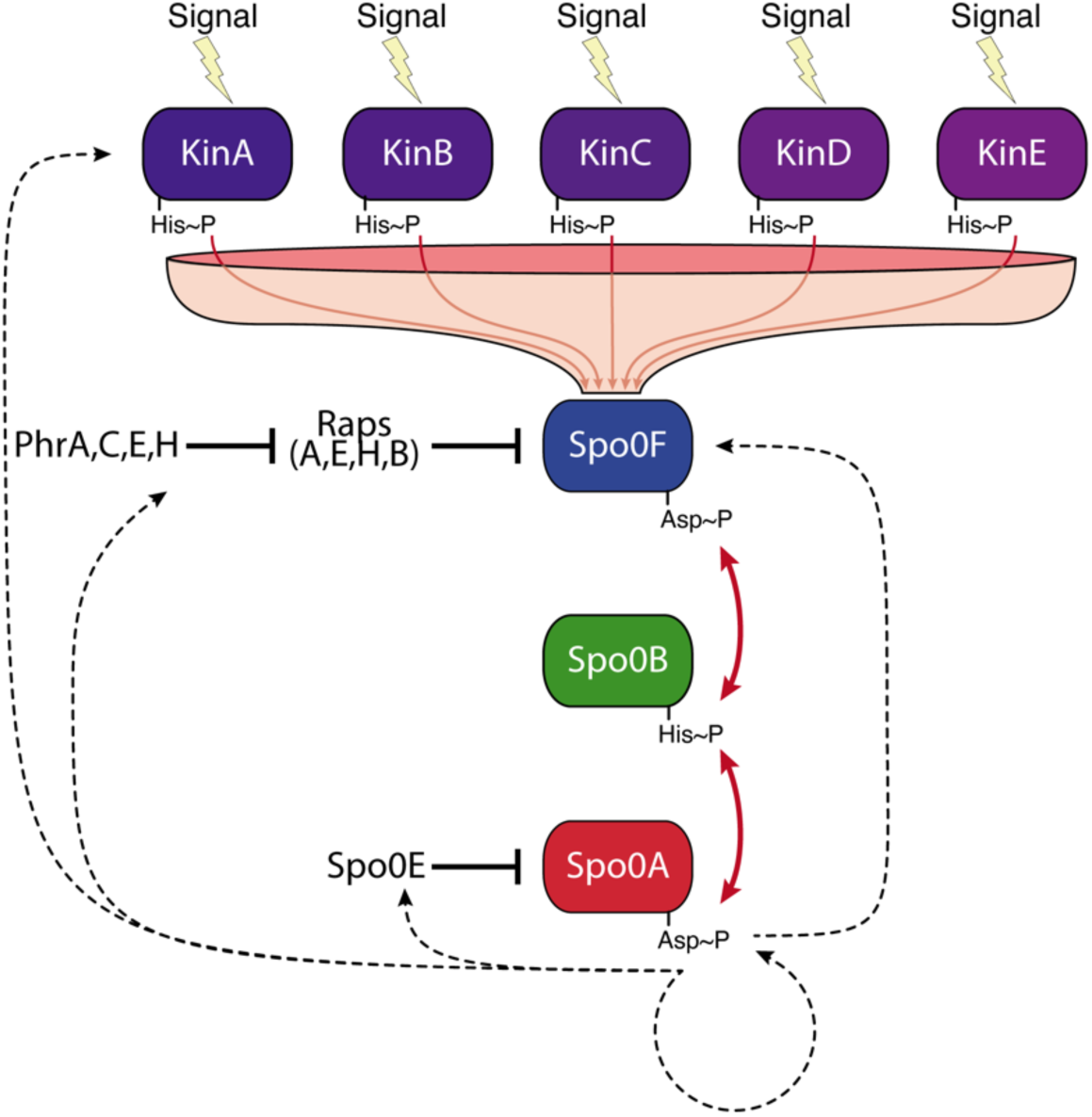
Sporulation phosphorelay pathway. Different signals are integrated through the action of multiple HKs (KinA to KinE) that funnel the information to the RR Spo0F. Information (in the form of a phosphoryl group) is then transmitted to the effector RR Spo0A through the phosphotransferase Spo0B. Dedicated Spo0F- and Spo0A-specific phosphatases are shown (Raps and Spo0E-like phosphatases, respectively). Red arrows indicate phosphoryl-transfer reactions. Dashed arrows represent the effect of transcriptional activation exerted by phosphorylated Spo0A on the other components of the pathway.

**Fig. S3:**
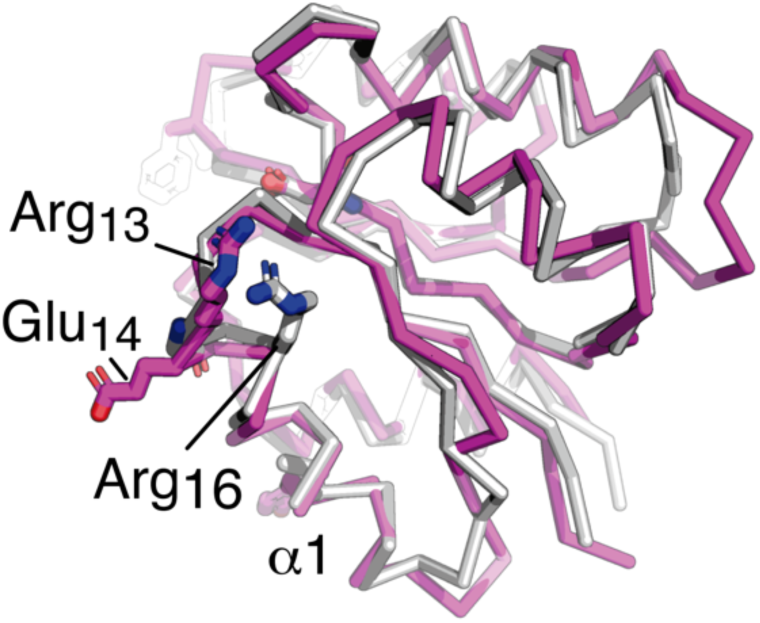
High structural similarity between Spo0F (from PDB Id 2FTK) and Spo0A.

**Fig. S4:**
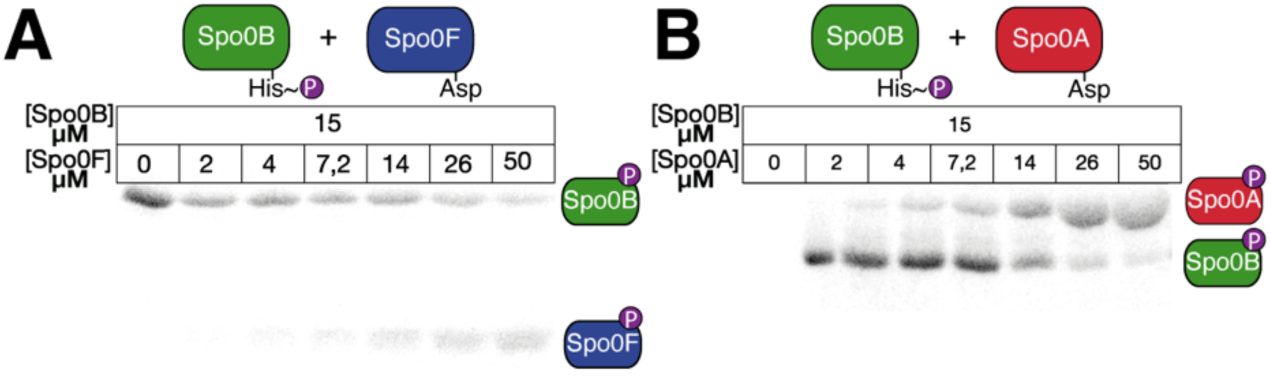
Phosphoryl-transfer equilibria in the B. subtilis sporulation phosphorelay pathway. A and B) Representative gel of a phosphotransfer titration assay between phosphorylated phosphotransferase Spo0B (P∼Spo0B) and the RRs Spo0F (A) or Spo0A (B) at equilibrium. Reactions were carried out with varying concentrations of RR from 0 to 50 µM at fixed level of Spo0B (15 µM) at pH 8 and 25 °C and analysed by SDS-PAGE and autoradiography.

**Fig. S5:**
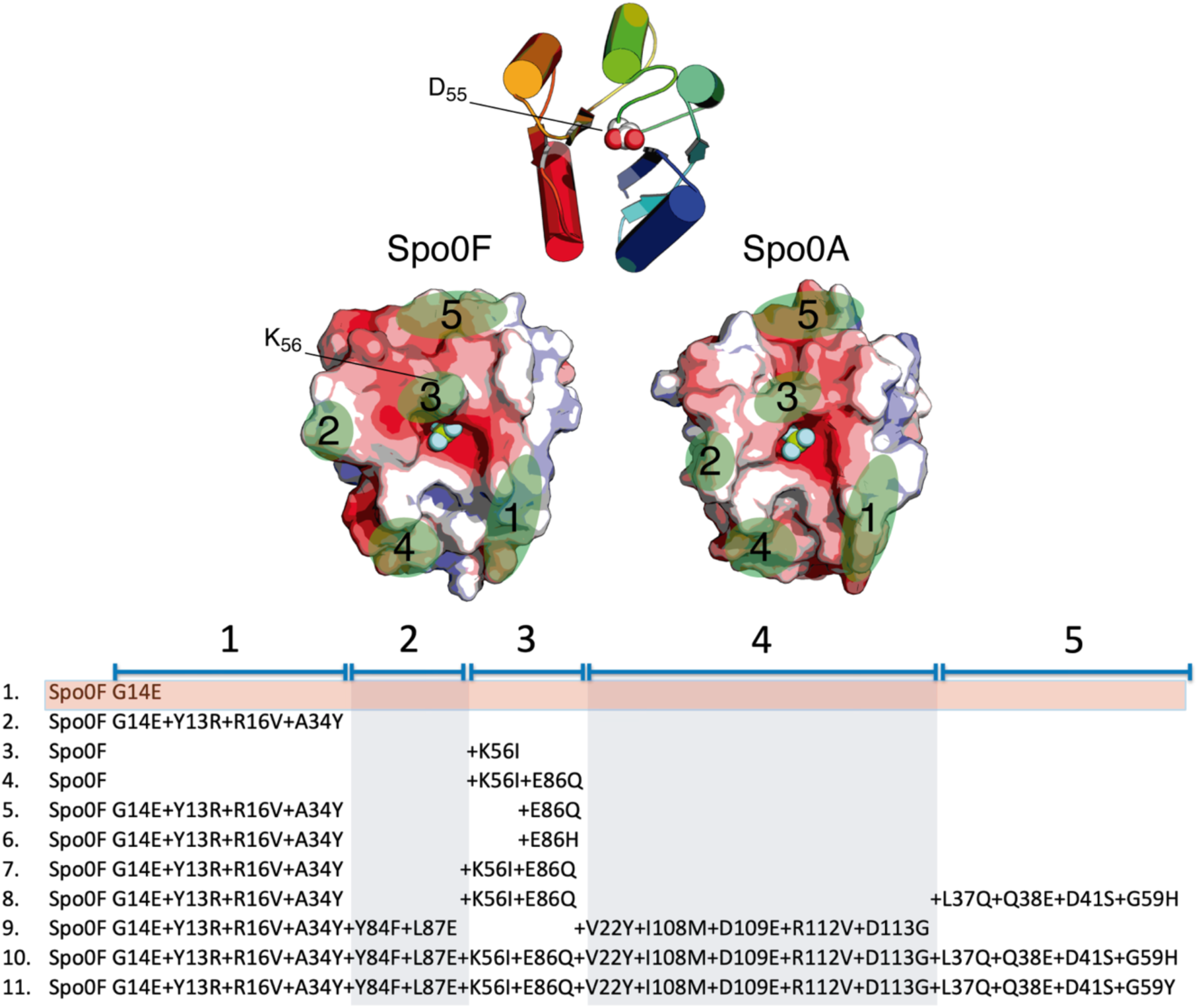
Grafting of motifs from Spo0A to Spo0F. A) View of the binding interface of Spo0F or Spo0A, to Spo0B. Spo0F and Spo0A are depicted in solvent-accessible surface representation. The surface is coloured according to the mapping of electrostatic potential (red=negative, blue=positive). Green patches indicate the selected sectors to graft motifs among different mutants. Table including all the substitutions included in Spo0F variants.

**Fig. S6:**
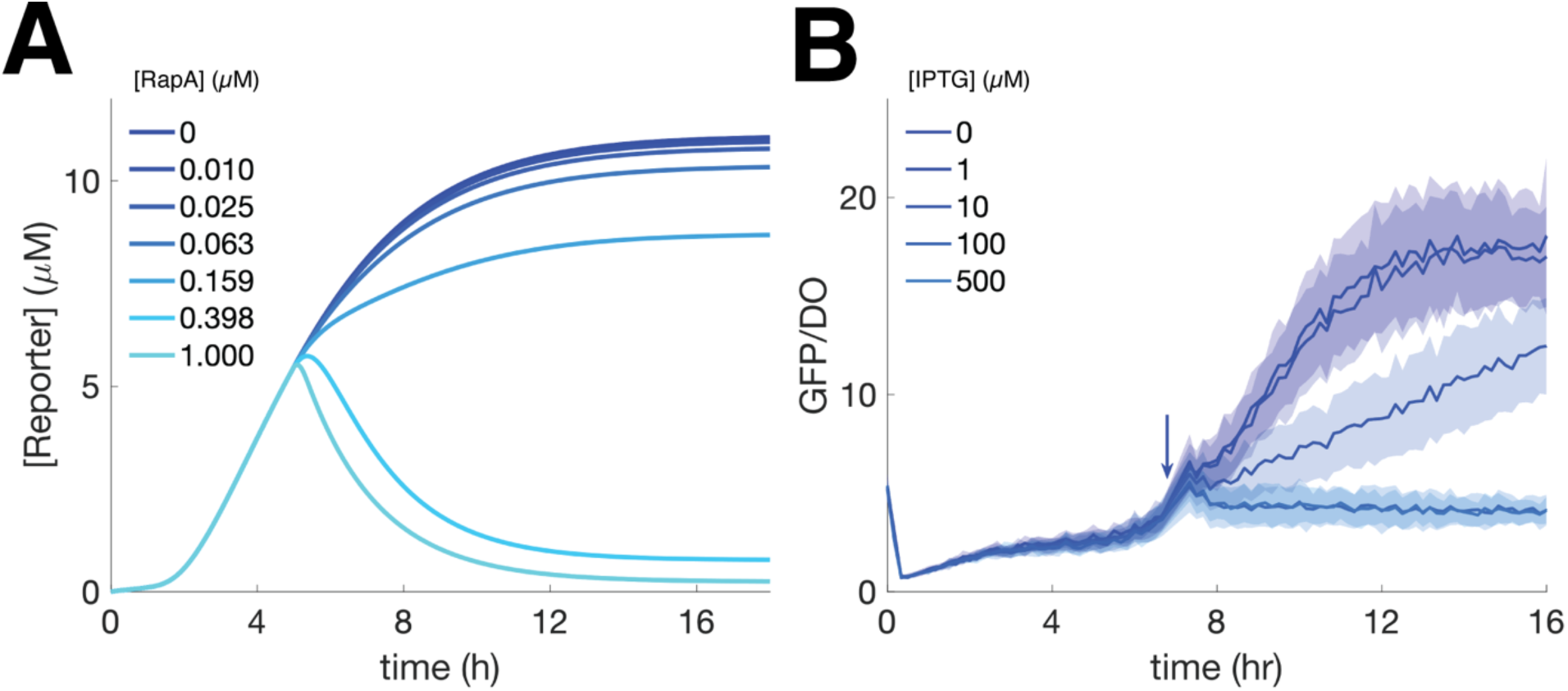
Resetting of the sporulation phosphorelay by phosphatase activity acting at the level of Spo0A. (A) Simulations of the sporulation system under activating conditions with wild type Spo0F, and under different steady state Spo0E concentrations, are shown. The reporter concentration is plotted as a function of time. (B) Time courses of sporulation in a *B. subtilis* strain expressing wild type Spo0F. Sporulation was induced by incubating cells in nutrient-deprived media under different IPTG concentrations (determining different Spo0E intracellular concentrations). Thick lines represent mean GFP/DO ratios, and the shaded region indicates one standard deviation

**Fig. S7.**
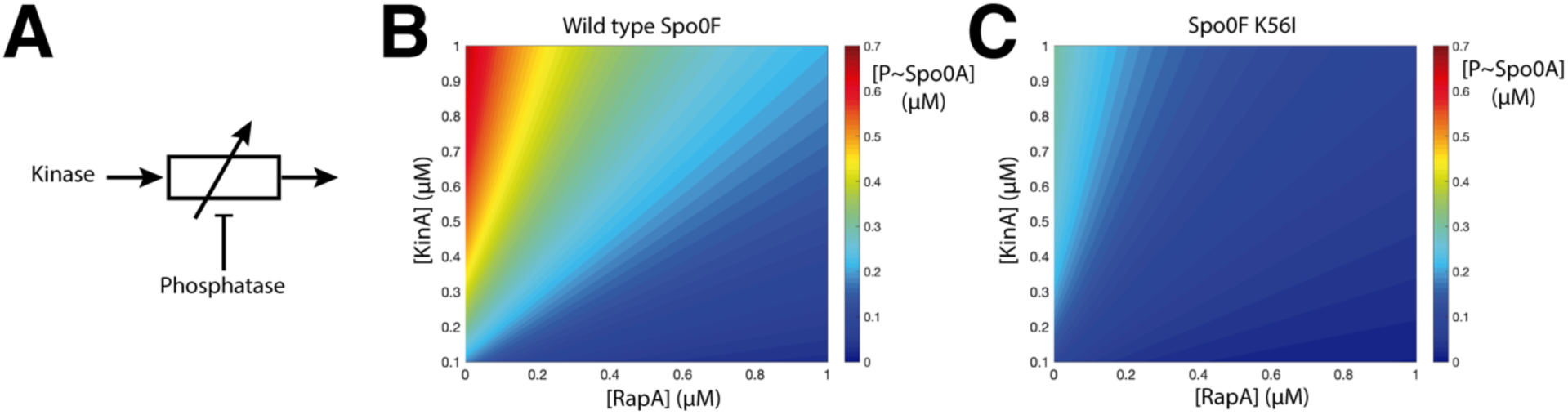
A Rheostat-like mechanism regulates Spo0A phosphorylation, leading to more graded responses. (A) Schematic representation of a rheostat-like regulatory mechanism for a signaling pathway. Simulations predicting the rate of Spo0A phosphorylation as a function of KinA and RapA concentrations, under unidirectional (B) or reversible (C) regimes. To visualize a clearer picture, these simulations were based on simplified models, where KinA was assumed to be constitutively active, and feedback loops regulating different sporulation components were not considered. The simulations were performed for 30 min. The colour ramp, from blue to red, indicates the concentration of phosphorylated Spo0A at the end of the simulation.

**Fig. S8:**
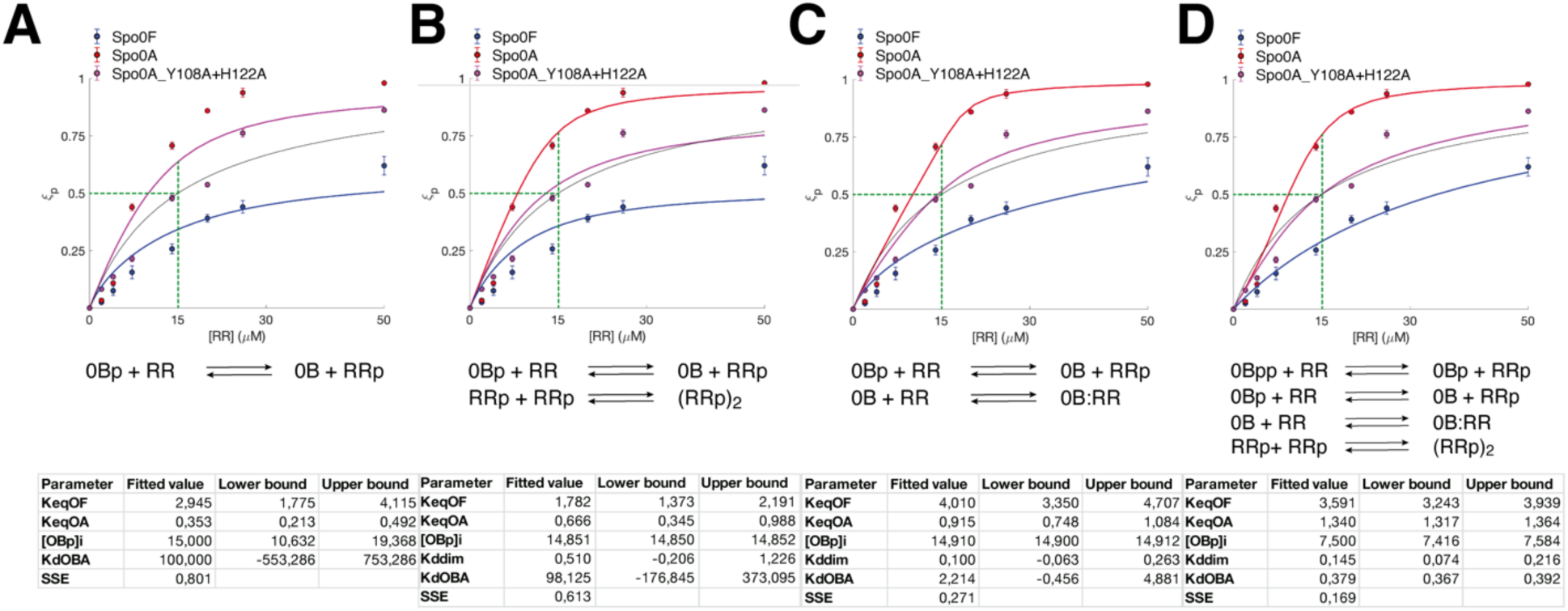
Fitting of kinetic models to phosphoryl-transfer titration experimental data. Each model was constructed as a custom script in MATLAB R2020a (MathWorks Inc.), consisting of a set of differential equations that describe each reaction. Continuous traces (red, magenta and blue lines) in (A) to (D) depict the prediction of the best-fitted model. The black continuous line shows the expected behaviour under a fully reversible equilibrium regime. Green lines indicate the reactions in equimolar concentrations.

**Table S1:** X ray diffraction data collection and refinement statistics.

| **Data collection** | Spo0B:Spo0A<sub>REC</sub> | Spo0B:Spo0A<sub>REC</sub> |
| --- | --- | --- |
| Space group | P2<sub>1</sub>2<sub>1</sub>2<sub>1</sub> | C2<sub>1</sub> |
| Cell dimensions |  |  |
| a, b, c (Å) | 76.361 125.504<br>149.472 | 221.240 103.145<br>66.461 |
| $\alpha, \beta, \gamma$ (°) | - | 90.0 96.91 90.0 |
| Wavelength (Å) | 0.97622 | 1.54179 |
| Resolution (Å) | 76.36 - 2.05 (2.08 -<br>2.05) | 28.72 - 3.50 (3.58-3.50) |
| Unique reflections | 90767 (4368) | 18675 (859) |
| R<sub>meas</sub> | 0.071 (1.506) | 0.143 (1.121) |
| R<sub>pim</sub> | 0.028 (0.667) | 0.073 (0.580) |
| CC<sub>1/2</sub> | 0.999 (0.466) | 0.998 (0.626) |
| I/σI | 11.15 (1.2) | 8.9 (1.5) |
| Completeness | 100 (97.7) | 91.3 (69.0) |
| Redundancy | 6.2 (5.0) | 3.8 (3.5) |
| Refinement |  |  |
| Resolution (Å) | 74.74 - 2.05 (2.06 -<br>2.05) | 24.02 - 3.50 (3.53 -<br>3.50) |
| Number of refls used<br>(N in the free set) | 4472 (85) | 915 (21) |
| R<sub>work</sub> / R<sub>free</sub> | 0.2235 / 0.2461 | 0.2426/0.292 |
| Number of atoms | 10209 | 6389 |
| Protein | 9749 | 6389 |
| Ligands + ions | 66 (GOL) | - |
| Water | 394 | - |
| B factors (Å<sup>2</sup>) |  |  |
| Wilson plot | 49.05 | 100.7 |
| Mean (overall) | 70.83 | 125.9 |
| R.m.s. deviations |  |  |
| Bond lengths (Å) | 0.008 | 0.010 |
| Bond angles (°) | 0.84 | 1.378 |
| Number of residues in<br>Ramachandran plot<sup>§</sup><br>(favored/allowed/outliers) | 1188/19/1 | 749/27/0 |
| PDB ID | pdb 00009PI0 | pdb 00009PHZ |

**Table S2:** Spo0F and Spo0A variant tested

|  |  |
| --- | --- |
| **F0** | Spo0F wild type |
| **F1** | Spo0F G14E |
| **F2** | Spo0F G14E+Y13R+R16V+A34Y |
| **F3** | Spo0F+K56I |
| **F4** | Spo0F+K56I+E86Q |
| **F5** | Spo0F G14E+Y13R+R16V+A34Y+E86Q |
| **F6** | Spo0F G14E+Y13R+R16V+A34Y+E86H |
| **F7** | Spo0F G14E+Y13R+R16V+A34Y+K56I+E86H |
| **F8** | Spo0F G14E+Y13R+R16V+A34Y+K56I+E86Q+L37Q+Q38E+D41S+G59H |
| **F9** | Spo0F G14E+Y13R+R16V+A34Y+Y84F+L87E+V22Y+I108M+D109E+R112V+D113G |
| **F10** | Spo0F<br>G14E+Y13R+R16V+A34Y+Y84F+L87E+K56I+E86Q+V22Y+I108M+D109E+R112V<br>+D113G+L37Q+Q38E+D41S+G59H |
| **F11** | Spo0F<br>G14E+Y13R+R16V+A34Y+Y84F+L87E+K56I+E86Q+V22Y+I108M+D109E+R112V<br>+D113G+L37Q+Q38E+D41S+G59Y |
| **A0** | Spo0A wild type |
| **A1** | Spo0A+Y108A+H122A |
| **A2** | Spo0A+E14G |
| **A3** | Receiver domain of Spo0A |

**Table S3.** Plasmids used in this work

| Name | Source |
| --- | --- |
| pMAD spo0E (EIB214) | This work |
| pMAD spo0F (EIB221) | Babel et al. |
| pMAD Spo0F-K56I | This work |
| pDR111 rapA (EIB297) | Mutlu et al. |
| pPspoiIE GFP BglS | This work |
| pRB117 Pspo0F mCherry | This work |
| pET-28b Spo0F F0 | This work |
| pET-28b Spo0F F1 | This work |
| pET-28b Spo0F F2 | This work |
| pET-28b Spo0F F3 | This work |
| pET-28b Spo0F F4 | This work |
| pET-28b Spo0F F5 | This work |
| pET-28b Spo0F F6 | This work |
| pET-28b Spo0F F8 | This work |
| pET-28b Spo0F F9 | This work |
| pET-28b Spo0F F10 | This work |
| pET-28b Spo0F F11 | This work |
| pQE80 Spo0A A0 | This work |
| pQE80 Spo0A A1 | This work |
| pQE80 Spo0A A2 | This work |
| pQE80 Spo0A A3 | This work |
| pQE80 Spo0A | This work |
| pETM KinA | This work |
| pQE80 Spo0B | This work |
| pET Spo0B strep | This work |

**Table S4.**
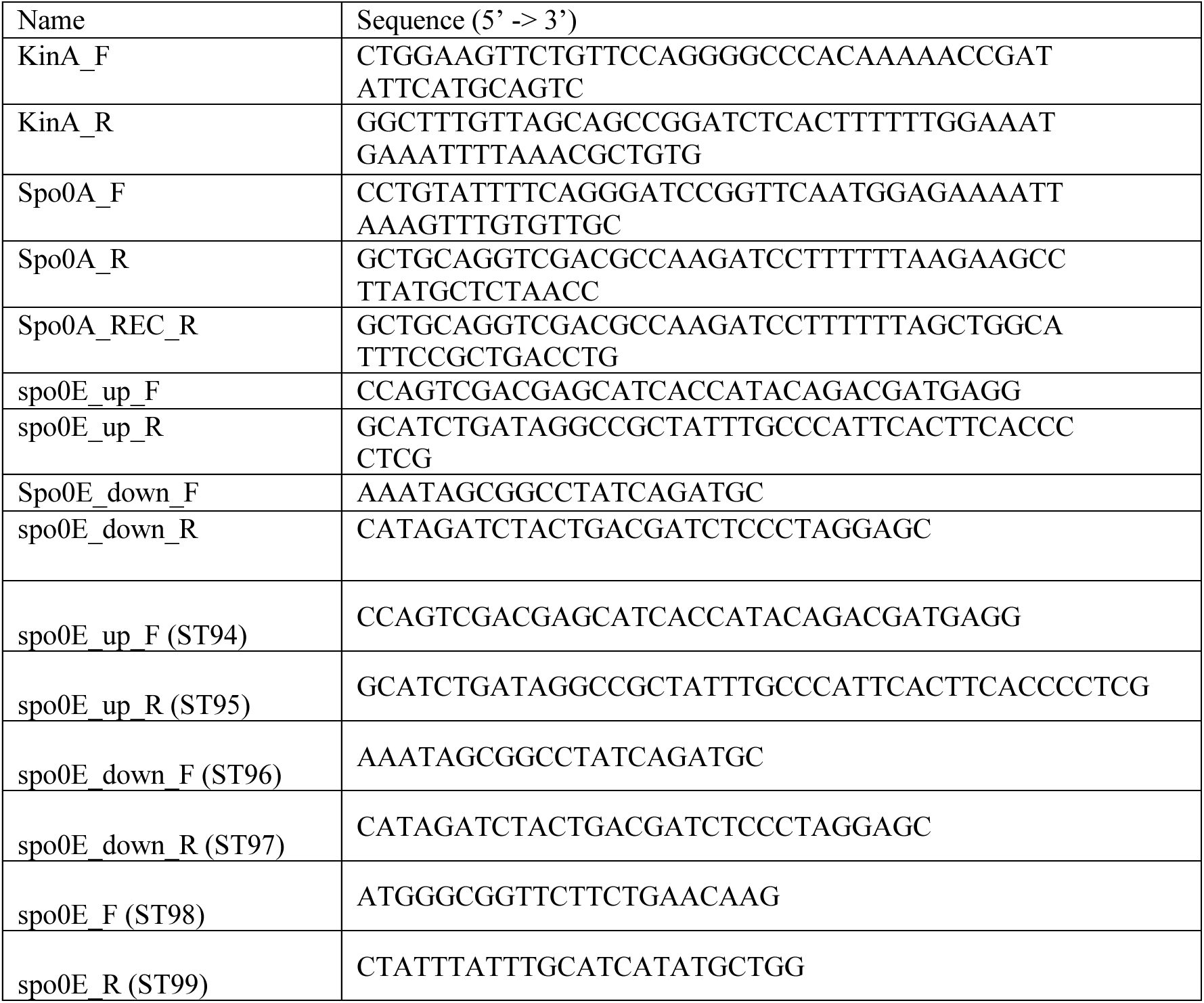
Primers used in this work

**Table S5.**
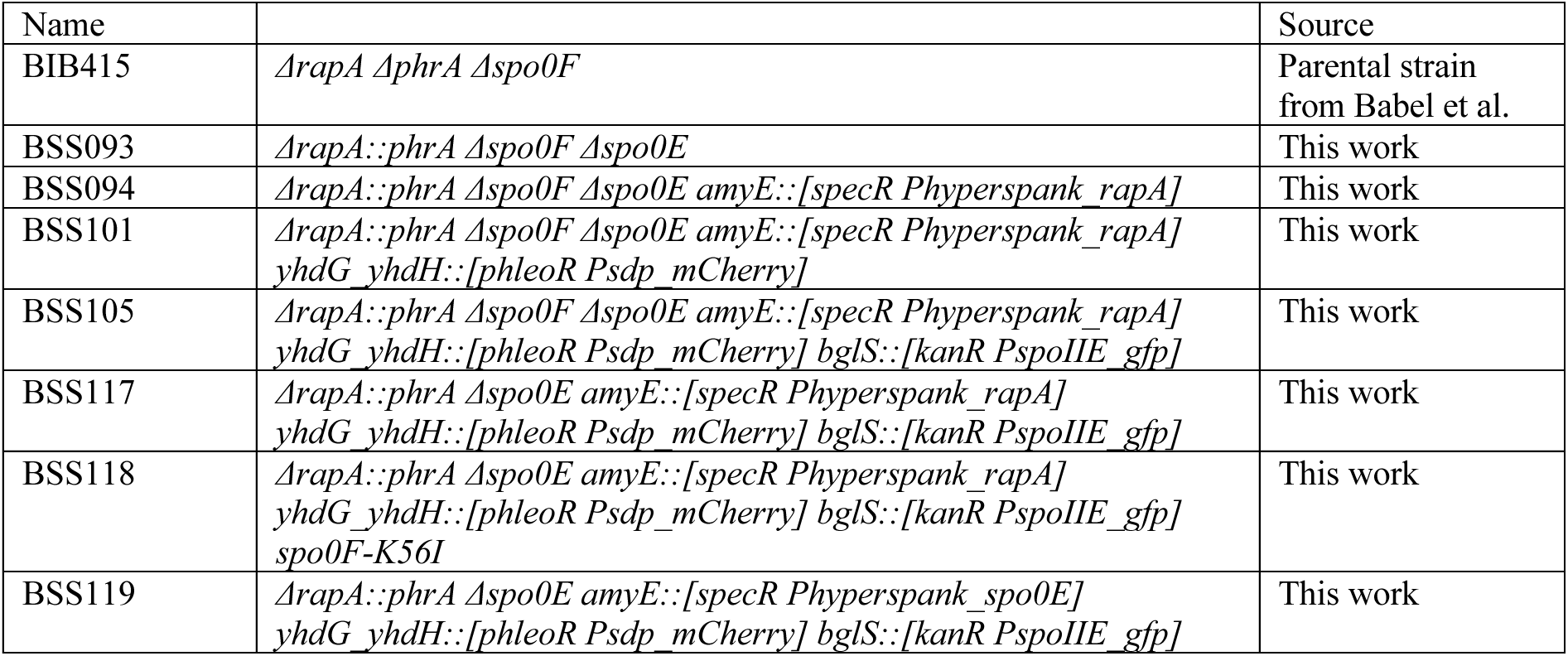
Strains used in this work

| Name |  | Source |
| --- | --- | --- |
| BIB415 | *ΔrapA ΔphrA Δspo0F* | Parental strain from Babel et al. |
| BSS093 | *ΔrapA::phrA Δspo0F Δspo0E* | This work |
| BSS094 | *ΔrapA::phrA Δspo0F Δspo0E amyE::[specR Phyperspank_rapA]* | This work |
| BSS101 | *ΔrapA::phrA Δspo0F Δspo0E amyE::[specR Phyperspank_rapA] yhdG yhdH::[phleoR Psdp_mCherry]* | This work |
| BSS105 | *ΔrapA::phrA Δspo0F Δspo0E amyE::[specR Phyperspank_rapA] yhdG yhdH::[phleoR Psdp_mCherry] bglS::[kanR PspoIIE_gfp]* | This work |
| BSS117 | *ΔrapA::phrA Δspo0E amyE::[specR Phyperspank_rapA] yhdG yhdH::[phleoR Psdp_mCherry] bglS::[kanR PspoIIE_gfp]* | This work |
| BSS118 | *ΔrapA::phrA Δspo0E amyE::[specR Phyperspank_rapA] yhdG yhdH::[phleoR Psdp_mCherry] bglS::[kanR PspoIIE_gfp] spo0F-K56I* | This work |
| BSS119 | *ΔrapA::phrA Δspo0E amyE::[specR Phyperspank_spo0E] yhdG yhdH::[phleoR Psdp_mCherry] bglS::[kanR PspoIIE_gfp]* | This work |

